# Genetic risk for schizophrenia influences substance use in emerging adulthood: An event-level polygenic prediction model

**DOI:** 10.1101/157636

**Authors:** Travis T. Mallard, K. Paige Harden, Kim Fromme

## Abstract

**Background:** Emerging adulthood is a peak period of risk for alcohol and illicit drug use. Recent advances in psychiatric genetics suggest that the co-occurrence of substance use and psychopathology arises, in part, from a shared genetic etiology. We sought to extend this research by investigating the influence of genetic risk for schizophrenia on trajectories of four substance use behaviors as they occurred across emerging adulthood.

**Method:** Young adult participants of non-Hispanic European descent provided DNA samples and completed daily reports of substance use for one month per year across four years (*N*=30,085 observations of *N*=342 participants). Polygenic scores for schizophrenia were included in two-level hierarchical linear models designed to test associations between genetic risk for schizophrenia, participant age, and four substance use phenotypes.

**Results:** Here, we interpret results at *p*<.05 as suggestive and results at *p*<.005 as significant. Accordingly, our results suggest that polygenic scores for schizophrenia were positively associated with participants’ overall likelihood to engage in illicit drug use, but not alcohol-related substance use. Moreover, our results indicate that participants with a greater polygenic loading for schizophrenia experienced greater age-related increases in the likelihood of using substances across emerging adulthood.

**Conclusions:** The present study used a novel combination of polygenic prediction and intensive longitudinal methods to characterize the influence of genetic risk for schizophrenia on patterns of age-related change in substance use across emerging adulthood. Results suggest that genetic risk for schizophrenia exerts both broad and developmentally-specific influences on substance use behaviors in a non-clinical population of young adults.

## Introduction

Emerging adulthood, which spans the ages of 18 to 25 years, is a peak developmental period for the initiation and escalation of alcohol and drug use (Kendler *et al*., 2008; Johnston *et al*., 2011). Approximately 75% of lifetime cases of substance use disorders develop by the mid-to late-20s (Christie *et al*., 1988; Kessler *et al*., 2005, 2007), and problematic substance use in this period often co-occurs with other forms of psychopathology (Grant *et al*., 2015, 2016). The presence of psychiatric comorbidity increases risk for negative health outcomes, contributing significantly to the morbidity and mortality associated with alcohol and drug use (Whiteford *et al*., 2013; Johnson *et al*., 2014). Whereas early research hypothesized that high rates of comorbidity reflected self-medication behaviors (i.e., efforts to alleviate distress engendered by schizophrenia symptoms), recent research has generated more compelling theories and models of comorbidity.

Advances in psychiatric genetics suggest that the co-occurrence of substance use and other mental health problems is due, in part, to a shared genetic etiology (Polimanti, Agrawal and Gelernter, 2017). That is, while a portion of the underlying genetic etiology of substance use may specifically increase liability for alcohol and/or drug use *per se*, other genetic risk factors for substance use may also be related to psychopathology more broadly (Johnson *et al*., 2009; Caspi *et al*., 2014; Pettersson, Larsson and Lichtenstein, 2016). Given the substantial heritability and polygenicity of substance use behaviors (Gratten *et al*., 2014; Polderman *et al*., 2015), it has been posited that some genetic variants dually confer risk for substance use *and* psychopathology, perhaps influencing biological pathways common to multiple psychiatric conditions (Cross-Disorder Group of the Psychiatric Genomics Consortium, 2013; Network and Pathway Analysis Subgroup of Psychiatric Genomics Consortium, 2015). Indeed, twin and family studies have reported that substance use behaviors arise from a heterogeneous etiology comprised of multiple genetic factors (Kendler *et al*., 2003, 2012).

Recently, polygenic scores have been used to examine shared, cross-trait genetic influences on several psychiatric phenotypes (Krapohl *et al*., 2016). Polygenic scores provide individual-specific estimates of genetic liability for a given trait by aggregating the effects of thousands of single nucleotide polymorphisms (SNPs) identified in large genome-wide association studies (GWASs). Because this approach leverages the results from well-powered GWASs, it is well-suited to the investigation of aggregate genetic effects with modest sample sizes (Belsky and Israel, 2014). Here, we apply this method in a university sample of emerging adults, where we examine the extent to which genetic risk for schizophrenia influences trajectories of alcohol and illicit drug use.

Our focus on genetic risk for schizophrenia is motivated by evidence suggesting that schizophrenia and substance use share a portion of their underlying genetic architecture (Polimanti, Agrawal and Gelernter, 2017). For instance, recent studies have found that schizophrenia shares modest but significant genetic correlations with cannabis use (Pasman *et al*., 2018), alcohol use (Clarke *et al*., 2017), and risk preferences (Linnér *et al*., 2018). Similarly, several cross-sectional studies have reported that polygenic scores for schizophrenia predict alcohol, amphetamine, cannabis, cocaine, opioid, and sedative use disorders (Power *et al*., 2014; Carey *et al*., 2016; Kalsi *et al*., 2016; Hartz *et al*., 2017; Reginsson *et al*., 2017; Verweij *et al*., 2017). However, while previous studies have related genetic risk for schizophrenia to diagnosed substance use disorders or lifetime substance use, no study has considered how this genetic risk functions in the context of development. That is, *when* does genetic risk for schizophrenia influence substance use?

The growing support for a shared genetic architecture between schizophrenia and substance use spans numerous studies, which employ various methodologies. In the present manuscript, we sought to extend this research through a person-centered intensive longitudinal approach that investigates the effect of genetic risk for schizophrenia on substance use as it occurred in the daily lives of emerging adults. To accomplish this aim, we first collected daily self-report data related to substance use from across a four-year period (*N*=30,085 observations, *M*=87.97 observations per person). We then extended polygenic prediction methods to event-level phenotypes, which increase measurement precision of behavior in the natural environment and can characterize within-person patterns of variation (Molenaar and Campbell, 2009). Finally, we constructed a hierarchical linear model (HLM) to test whether genetic risk shapes how substance use *changes* across emerging adulthood—a developmental period in which genetic risk for schizophrenia often manifests (Kessler *et al*., 2007).

To calculate polygenic scores for schizophrenia, we used results from the Psychiatric Genomic Consortium’s (PGC) most recent GWAS of the disorder (Schizophrenia Working Group of the Psychiatric Genomics Consortium, 2014). We then investigated the effect of genetic risk for schizophrenia on four event-level phenotypes: daily alcohol use, binge drinking, illicit drug use, and concurrent alcohol and drug use. Specifically, we tested: (i) whether polygenic scores for schizophrenia predicted an individual’s overall likelihood to engage in substance use on a given day, and (ii) whether polygenic scores for schizophrenia predicted the magnitude of longitudinal, age-related change in substance use. In accordance with previous research, we hypothesized that genetic risk for schizophrenia would be positively associated with all forms of substance use. Furthermore, given that schizophrenia often onsets between late adolescence and early adulthood, we hypothesized that genetic risk for schizophrenia would be associated with a greater likelihood to use substances as participants grow older. In testing these hypotheses, we hope to lend insight into the heterogeneous genetic etiology of substance use behaviors and when they manifest in development.

## Materials and Methods

### Participants

The present sample was recruited from a larger cohort of subjects who participated in a longitudinal investigation of alcohol abuse and behavioral risks among college students. Recruitment procedures for the full study have been described in previously published articles (Fromme, Corbin and Kruse, 2008; Ashenhurst *et al*., 2015; Mallard *et al.*, 2018). A subset of the full sample completed a daily monitoring protocol and provided DNA for genotyping procedures (*n*=541, 64% non-Hispanic European, 67% female). To avoid potential effects associated with population stratification, the analyses detailed below were limited to the non-Hispanic European portion of the sample (*n*=354, 66% female). Twelve participants were excluded from analyses following the quality control procedures described below (Final *N*=342, 66% female, *M*_age_=18.44 years, *SD*_age_=0.32 years). The university’s Institutional Review Board approved all study procedures.

### Genotyping protocol and quality control

Participants provided 2 mL of saliva in Oragene-Discover (Oragene™, DNAgenotek, Ottawa, Ontario, Canada) collection kits that were distributed and returned via mail. DNA for participants was extracted, purified, and diluted from saliva samples at the Institute for Behavior Genetics at the University of Colorado, Boulder. Purified and diluted samples were sent to the Neuroscience Genomics Core at the University of California, Los Angeles, for chip-based genotyping. Samples were assayed on an Illumina BeadLab platform using an Illumina Infinium PsychArray BeadChip array (San Diego, CA), which assays ∼265,000 SNPs across the genome.

Genotypic data were subjected to quality control procedures recommended for chip-based genomic data (Anderson *et al*., 2010; Turner *et al*., 2011). Samples were excluded from statistical analyses because of poor call rate (<98%), inconsistent self-reported sex and biological sex, and relatedness (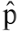 >.125). SNPs were excluded from analyses if more than 2% of genotype data was missing. Thresholds for minor allele frequency (MAF) and Hardy-Weinberg Equilibrium (HWE) were applied after phasing and imputation (described below), as variant-level filtering has been shown to have a detrimental effect on imputation quality (Roshyara *et al*., 2014).

Finally, although the present analyses were limited to participants who self-reported non-Hispanic European descent, flashPCA2 was used to (i) extract the top ten genomic principal components of ancestry and (ii) identify ancestral outliers (i.e., participants with a greater level of admixture than reported). First, principal components of ancestry were estimated using the European samples from Phase 3 v5 of the 1000 Genomes Project (1000 Genomes Project Consortium *et al*., 2015) as a reference sample. Outliers were then defined as any participant with a score greater than or equal to four standard deviations from the mean on the first and/or second principal component of ancestry (i.e., the range present in European samples from Phase 3 v5 of the 1000 Genomes Project); five participants met this exclusion criterion. Scatterplots of the principal component scores were then examined to confirm that no ancestral outliers remained in the sample.

### Imputation

Unknown genotypes were imputed on the *Michigan Imputation Server* (https://imputationserver.sph.umich.edu). Variants were phased with Eagle v2.3 (Loh, Palamara and Price, 2016) and imputed with Minimac3 1.0.13 (Das *et al*., 2016), using Phase 3 v5 of the 1000 Genomes Project (1000 Genomes Project Consortium *et al*., 2015) as a reference panel. To ensure all markers were of high quality, several post-imputation quality control thresholds were applied. After phasing and imputation, SNPs with a MAF < .01, INFO score < .90, or HWE *P*-value < .00001 were excluded from all statistical analyses. These procedures yielded a final set of 5,250,123 high-quality genotyped and imputed variants.

### Genome-wide polygenic scores

Polygenic scores for schizophrenia were calculated for 342 unrelated participants of non-Hispanic European ancestry by using summary statistics from the PGC’s 2014 GWAS of schizophrenia (Schizophrenia Working Group of the Psychiatric Genomics Consortium, 2014). Specifically, summary statistics were obtained for the 15,358,497 variants analyzed in PGC cohorts of European ancestry, which consisted of 32,405 cases, 42,221 controls, and 1,235 trios. These variants were restricted to 4,509,191 bi-allelic SNPs that were present in both datasets after the quality control procedures described above. LD-based clumping was then used to identify a set of 121,702 independent SNPs (*r*^2^ < 0.1 in the present sample) with the lowest *p*-value in a given 1 Mb window. An additional LD threshold was imposed to ensure that these independent SNPs were not in long-range LD with each other (*r*^2^ > 0.1 within a 10 Mb window). This process identified a final set of 118,719 independent SNPs to be used for the polygenic scores.

Before calculating polygenic scores, the odds ratios reported by the PGC were log-transformed to identify the beta coefficients associated with the effect allele for each SNP. PLINK 1.9 (Chang *et al*., 2015) was then used to calculate a polygenic score for each participant by multiplying the number of effect alleles (0, 1, or 2) at a given SNP by its associated beta coefficient and summing across all included SNPs. Finally, polygenic scores were *z*-standardized to aid interpretation of results, establishing a mean (*M*) of 0 and a standard deviation (*SD*) of 1 (Figure 1).

**Figure 1.**
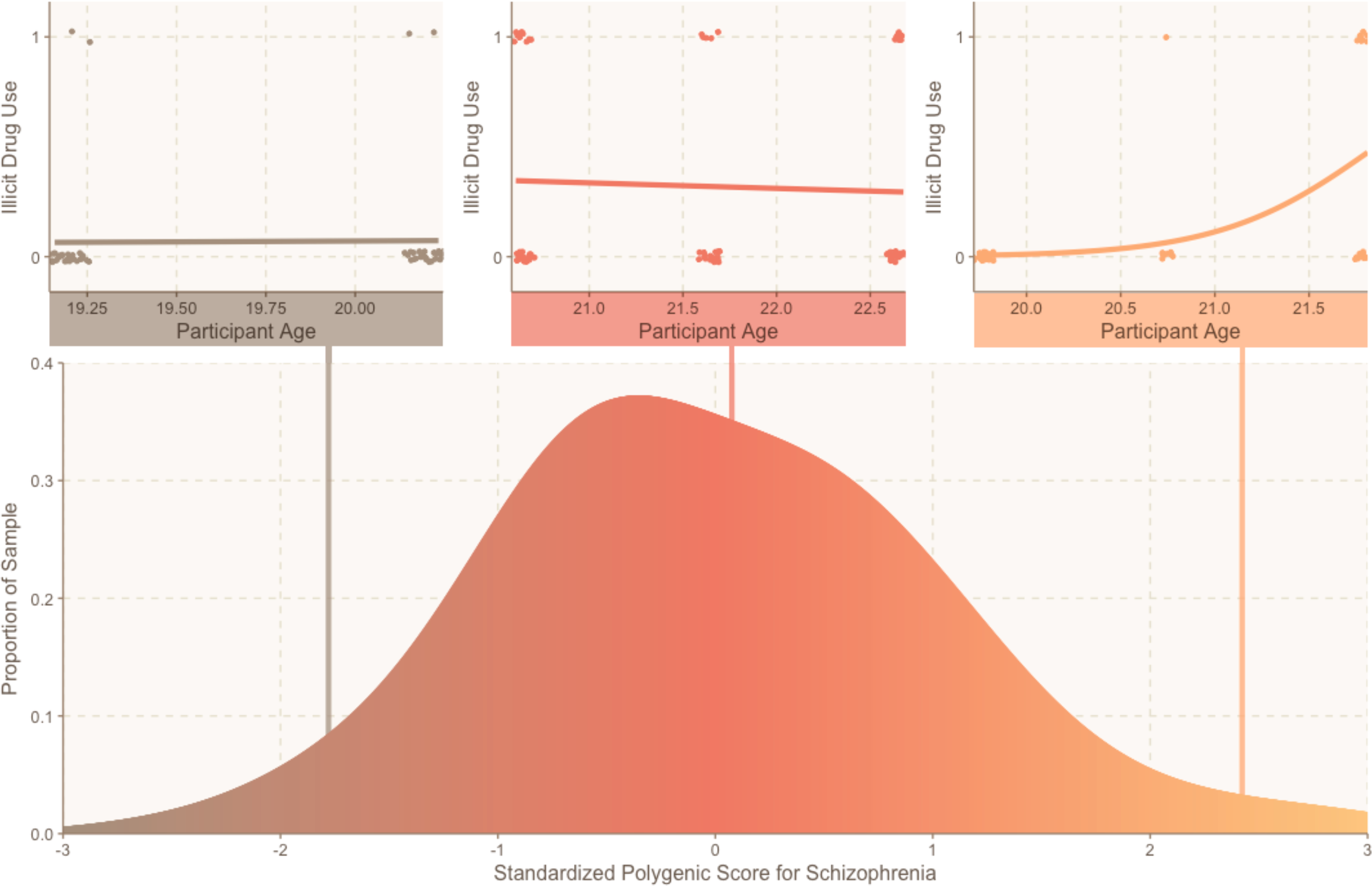
A density plot illustrating the distribution of standardized polygenic scores for schizophrenia in the present sample. The top three panels show data from three randomly selected participants who have different polygenic scores for schizophrenia. Each dot represents one event-level observation during which illicit drug use did occur (plotted as a “1” on the Y-axis) or did not occur (plotted as a “0” on the Y-axis). The X-axis of the panel represents the age of the participant at each event-level observation. The line represents the person-specific line of best fit predicting probability of illicit drug use.

### Longitudinal event-level design and phenotyping

Participants completed up to 30 consecutive days of online self-monitoring in each of their first four years of college. At the beginning of the study, a random sample of 200 students were invited to participate in a daily monitoring study. A random selection of 40-43 students thereafter were invited to participate in the study each week to ensure sufficient monitoring across the entire calendar year. During their annual reporting period, participants were instructed to use the self-monitoring website (maintained by DatStat, Seattle, WA) to answer questions about the previous day.

Each day, participants answered questions about the previous day related to time-varying characteristics (e.g., weight), alcohol consumption (“*How many drinks did you consume yesterday?” and “Of the times that you drank this day, how long was your heaviest drinking episode?*”), and illicit drug use (“*Did you use illicit drugs yesterday?*”). If participants endorsed illicit drug use on any given day, they were asked to specify whether the drug use occurred while sober or during a drinking episode. Four event-level substance use phenotypes were assessed: any alcohol use, binge drinking, illicit drug use, and concurrent alcohol and drug use. Operant definitions for these substance use phenotypes are presented below.

- Alcohol use was defined as any consuming at least one standard drink during the reporting day.
- Binge drinking was defined as consuming alcohol at a rate of 2 or 2.5 standard drinks per hour for at least two hours (i.e., equivalent to the NIAAA definition of 4 or 5 drinks within a 2-hour period, depending on sex).
- Illicit drug use was defined as consumption of any illicit drugs during the reporting day.
- Concurrent alcohol and drug use was defined as any simultaneous consumption of alcohol and drugs during the reporting day.

Additionally, the self-monitoring website recorded the time and date of each daily report, which was used to determine the participant’s age (rounded to two decimal points) on a given day. This approach allowed us to model age as a continuous event-level predictor that varied within a 30-day reporting period (e.g., increasing 17.92 to 18.05 during the first reporting period), as well as between reporting periods (e.g., increasing from 18.05 to 18.92 between the first and second reporting periods). To reduce potential bias attributable to over-exclusion or inclusion of noncompliant participants, eight participants who did not provide at least 14 days of monitoring data were excluded from statistical analyses. The final sample included 342 participants with 30,085 event-level observations.

### Analytic approach

A two-level hierarchical linear model (HLM; Raudenbush and Bryk, 2002) with robust standard errors was used to analyze the relationships between polygenic scores for schizophrenia (PGS_SCZ_), participant age (AGE), and the four substance use phenotypes. Events were nested within participants for all statistical analyses. As between-person and within-person relationships are not necessarily synonymous (Figure 1; Molenaar and Campbell, 2009), the HLM included a random intercept and random slope to account for individual differences in the overall level of substance use and rate of age-related change in substance use, respectively. Principal components of ancestry (PC_1_…PC_10_), biological sex (SEX), and age at beginning of college (AGE_W1_) were included as trait-level covariates in all analyses. The full model is described below.

#### LEVEL 1 MODEL

Prob(OUTCOME = 1|π) = φ

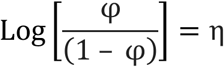

η =π_0_ + π_1_ (AGE)

#### LEVEL 2 MODEL

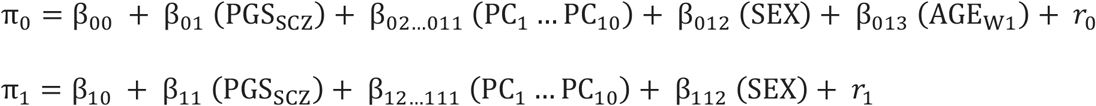

All substance use phenotypes were analyzed using a logit model. The Level 1 (event level) equation modeled the likelihood of a participant engaging in substance use on a given day as a function of a person-specific random intercept (π_0_) and a person-centered random slope describing within-person variability in the likelihood of using substances as a function of event-level age (π_1_). Importantly, event-level age was centered on the person mean and thus reflects within-person, age-related change in substance use over time (Raudenbush and Bryk, 2002). Overall, the Level 1 equation tested the extent to which a person showed systematic age-related change in substance use.

The Level 2 (person level) equation modeled between-person variability in the likelihood to use substances when aggregating across all occasions. Here, the intercept for substance use phenotypes (π_0_), which represents the person-average likelihood to engage in substance use across all events, was modeled as a function of the effect of polygenic scores for schizophrenia (β_01_), as well as the effects of ancestry (β_02_…β_11_), sex (β_12_), and age at first wave of data collection (β_13_). We additionally modeled the random slopes for event-level age as a function of the effects of polygenic scores for schizophrenia (β_11_), ancestry (β_12_…β_111_), and sex (β_112_). The first ten principal components of ancestry and age at first wave were centered on the grand mean, while polygenic scores for schizophrenia and sex were uncentered. Between-person residuals were included for all event-level slopes (*r*_0_ and *r*_1_) to allow for heterogeneity in the magnitude of within-person effects. Overall, the Level 2 model tested whether polygenic scores for schizophrenia, sex, and mean age predicted (i) participants’ overall likelihood to use substances when aggregating across all events and (ii) age-related changes in the likelihood to use substances as participants grew older.

## Results

Per recent reflections on statistical power and reproducibility (Benjamin *et al*., 2018), we interpret results at *p* < .05 as suggestive and results at *p* < .005 as significant. Descriptive statistics for each substance use phenotype are presented stratified by year in Table 1. Throughout the course of the study, alcohol consumption and binge drinking were reported at least once by 92.4% and 74.3% of the sample, respectively, while illicit drug use and concurrent alcohol and drug use were reported by 30.1% and 23.1% of the sample, respectively. The number of reporting days that each participant completed was not associated with age or polygenic scores for schizophrenia, though male sex was negatively associated with the number of daily monitoring reports completed (*r* = −.183, *p* = .001).

**Table 1.** Descriptive statistics for each substance use phenotype stratified by reporting year

|  | Year 1 (n = 317) |  | Year 1 (n = 321) |  | Year 1 (n = 309) |  | Year 1 (n = 236) |  | Total (N = 342) |  |
| --- | --- | --- | --- | --- | --- | --- | --- | --- | --- | --- |
|  | Non-Abstainers | Events | Non-Abstainers | Events | Non-Abstainers | Events | Non-Abstainers | Events | Non-Abstainers | Events |
| Alcohol Use | 198 | 998 | 220 | 1326 | 246 | 1824 | 205 | 1522 | 316 | 5670 |
| Binge Drinking | 120 | 371 | 145 | 504 | 163 | 591 | 142 | 434 | 254 | 1900 |
| Illicit Drug Use | 51 | 163 | 57 | 329 | 48 | 292 | 33 | 192 | 103 | 976 |
| Concurrent Alcohol and Drug Use | 28 | 68 | 43 | 132 | 35 | 112 | 29 | 85 | 79 | 397 |

The effects of polygenic scores for schizophrenia on the intercept and slope of all four substance use phenotypes are presented in Figure 2 and Table 2. Here, we represent the effects of polygenic scores for schizophrenia as odds ratios, which reflect change in the odds of an outcome given a one unit increase in the predictor. Moreover, as we standardized all polygenic scores prior to analysis, we characterize how a 1 *SD* increase in genetic risk for schizophrenia influences the likelihood to use substances on any given reporting day.

**Figure 2.**
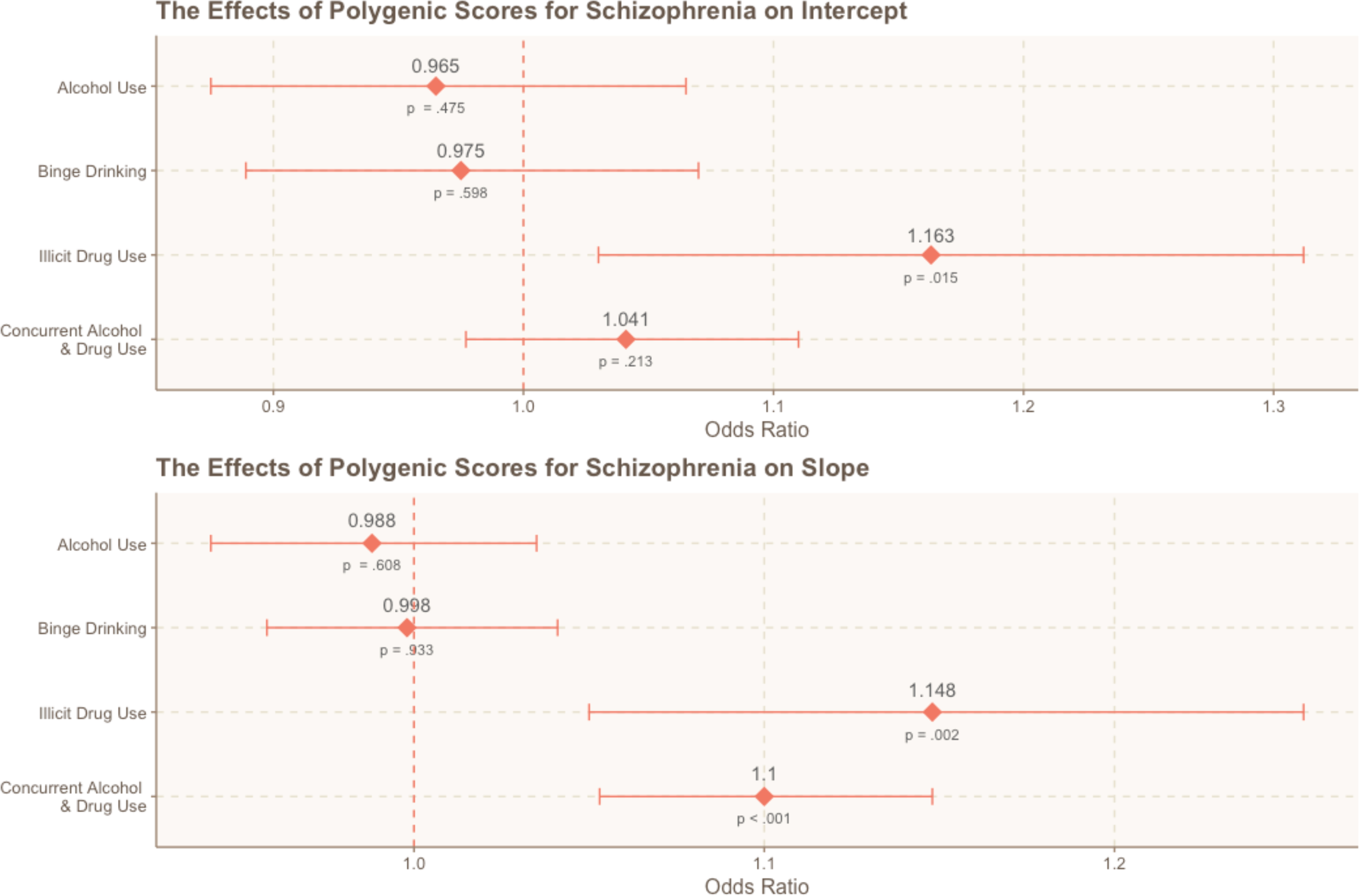
Odds ratios illustrating the effects of genetic risk for schizophrenia on the intercept and age-related slope of the five substance use phenotypes. The bars for each estimate reflect the 95% confidence interval, while the corresponding p-value is listed below each point.

**Table 2.** The effect of polygenic scores for schizophrenia on the intercept and slope of each substance use phenotype

| Intercept | *B* | OR | 95% CI | *p*-value |
| --- | --- | --- | --- | --- |
| Alcohol Use | -.036 | .965 | (.875,1.065) | .475 |
| Binge Drinking | -.025 | .975 | (.889,1.070) | .598 |
| Illicit Drug Use | .151 | 1.163 | (1.030,1.312) | .015<sup>†</sup> |
| Concurrent Alcohol and Drug Use | .040 | 1.041 | (0.977,1.110) | .213 |
| Slope | *B* | OR | 95% CI | *p*-value |
| Alcohol Use | -.012 | .988 | (.942,1.035) | .608 |
| Binge Drinking | -.002 | .998 | (.958,1.041) | .933 |
| Illicit Drug Use | .138 | 1.148 | (1.050,1.254) | .002<sup>*</sup> |
| Concurrent Alcohol and Drug Use | .095 | 1.100 | (1.053,1.148) | <.001<sup>*</sup> |

Note: OR = Odds Ratio, CI = Confidence Interval. <sup>†</sup> = suggestive of statistical significance at $p < .05$ . <sup>\*</sup> = statistically significant at $p < .005$ (Benjamin *et al.*, 2018).

We observed significant effects of polygenic scores for schizophrenia on illicit drug use and concurrent alcohol use and drug use. However, polygenic scores for schizophrenia did not influence the likelihood to engage in any alcohol use or binge drinking. Additionally, biological sex was not associated with any form of substance use, but age at the beginning of college (i.e., between-person differences in age) was suggestively associated with all types of use (all *p* < .05. Getting older over the course of the study (i.e., event-level age) was also positively associated with a greater likelihood to engage in all forms of substance use (all *p* < .001; Figure 3). The specific results for the drug use and concurrent alcohol and drug use phenotypes are detailed below.

**Figure 3.**
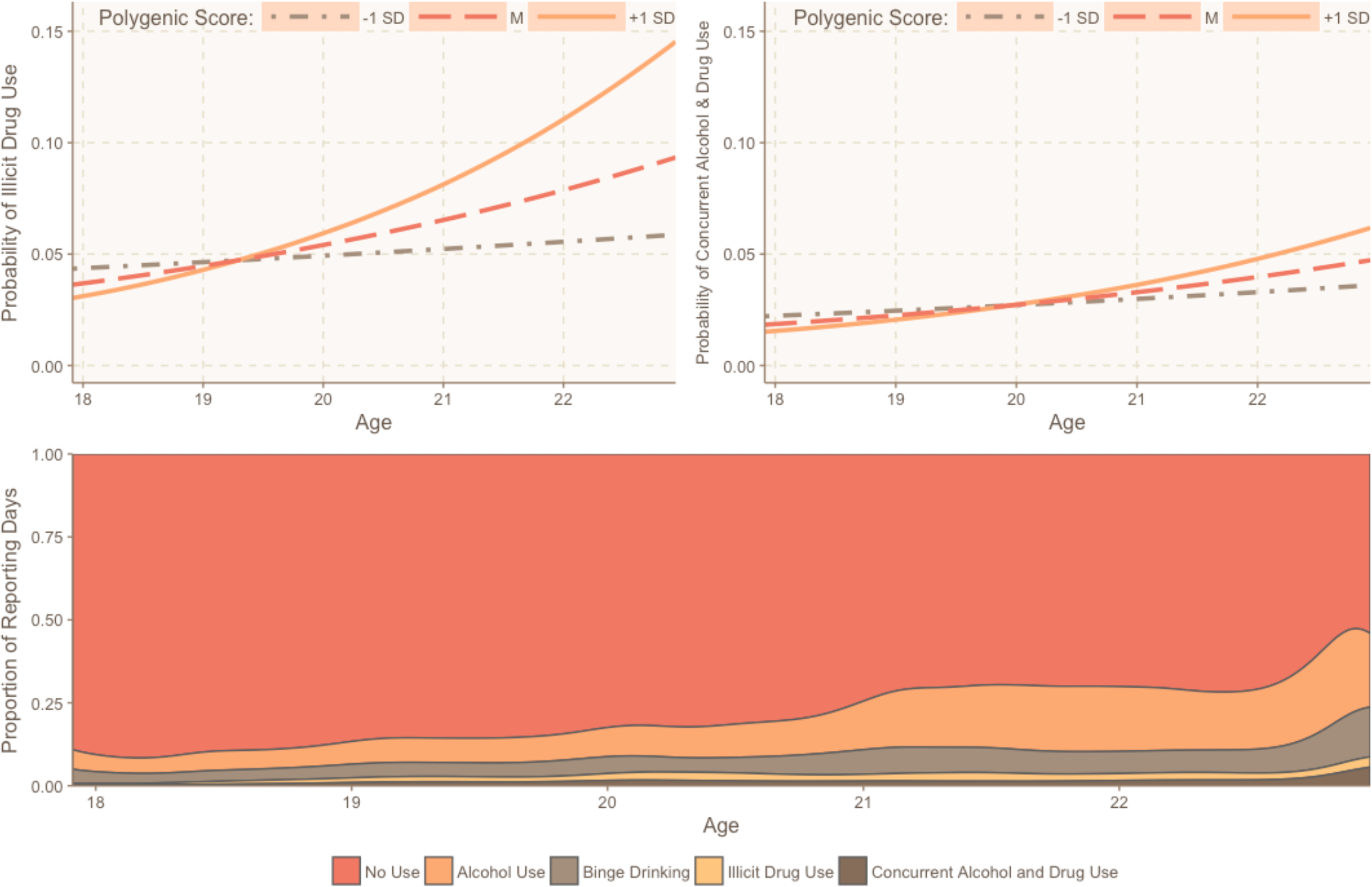
Above, the effects of genetic risk for schizophrenia on the likelihood to engage in illicit drug use and concurrent alcohol and drug use, as measured in the present study. In both cases, we see that greater genetic risk for schizophrenia predicts a more substantial increase in age-related substance use. Below, a scaled density plot illustrates the proportion of reporting days that included substance use in the present study.

### Illicit drug use

Results suggested that polygenic scores for schizophrenia predicted a greater overall likelihood to use illicit drugs (*B* = .151, OR = 1.163, *p* = .015). Here, a 1 *SD* increase in genetic risk for schizophrenia was associated with a relative 16.3% increase in the likelihood to engage in illicit drug use on any given day (i.e., across the entire study). Furthermore, results indicated that age-related changes in illicit drug use varied as a function of polygenic scores for schizophrenia. Specifically, we found that genetic risk for schizophrenia was significantly associated with the event-level slope between age and illicit drug use (*B* = .138, OR = 1.148, *p* = .002). So, participants with higher polygenic scores for schizophrenia were more likely to use illicit substances overall *and* experienced a more substantial increase in the likelihood of using drugs as they grew older. This effect is illustrated in Figure 3.

### Concurrent alcohol and drug use

Polygenic scores for schizophrenia did not predict a greater overall likelihood to engage in concurrent alcohol and drug use (*B* = .040, OR = 1.041, *p* > .05). However, results indicated that genetic risk for schizophrenia did significantly influence age-related change in concurrent alcohol and drug use. Specifically, we found that polygenic scores for schizophrenia were positively associated with the event-level slope between age and illicit drug use (*B* = .095, OR = 1.010, *p* < .001), such that participants with higher polygenic scores for schizophrenia experienced a more substantial increase in the likelihood of concurrent alcohol and drug use as they grew older. This effect is illustrated in Figure 3.

### Discussion

The present paper describes the first longitudinal, event-level examination of genetic risk for schizophrenia and its effects on daily substance use in a sample of university students. Specifically, we tested whether (i) polygenic scores for schizophrenia predicted a greater overall likelihood to engage in substance use on a given day and (ii) within-person age-related changes in substance use varied as a function of polygenic scores for schizophrenia. We report two major findings. First, we found suggestive evidence that genetic risk for schizophrenia predicted an individual’s overall likelihood to engage in illicit drug use, but it did not predict the likelihood that participants would engage in any form of alcohol-related substance use. Second, we found that genetic risk for schizophrenia significantly predicted the rate of age-related change in illicit drug use and concurrent alcohol and drug use.

Overall, a greater polygenic loading for schizophrenia predicted more recurrent illicit drug use. Whereas many prior studies have only examined the effect of polygenic scores for schizophrenia on substance use with diagnostic phenotypes or lifetime use outcomes, we identified genetic influences on substance use in the daily lives of emerging adults. Moreover, we found that participants with a greater polygenic loading for schizophrenia experienced greater age-related increases in the likelihood of using substances across emerging adulthood. As a result, our findings corroborate and build upon recent studies that have reported associations between polygenic scores for schizophrenia and problematic substance use (Power *et al*., 2014; Carey *et al*., 2016; Kalsi *et al*., 2016; Reginsson *et al*., 2017; Verweij *et al*., 2017).

Although our daily measure of illicit drug use did not identify the specific substance that was consumed, related investigations of this cohort have identified cannabis as the most commonly used illicit drug (Fromme, Corbin and Kruse, 2008). As previously noted, schizophrenia and cannabis use share a modest but significant genetic correlation (*r_g_* = .24; Pasman *et al*., 2018). Researchers have recently begun to interrogate the relationship between these two phenotypes, finding that genetic risk for schizophrenia exerts a causal influence on the liability to use cannabis (Pasman *et al*., 2018). One possibility is that genetic variants that confer risk for schizophrenia also influence cannabis use by impacting some shared pathophysiology (Chambers, Krystal and Self, 2001; Cross-Disorder Group of the Psychiatric Genomics Consortium, 2013; Network and Pathway Analysis Subgroup of Psychiatric Genomics Consortium, 2015). Alternatively, individuals with a higher polygenic loading for schizophrenia may experience prodromal symptoms or neurocognitive impairment that leads them to use cannabis (i.e., a graded iteration of the “self-medication” hypothesis). Our results indicate that genetic risk for schizophrenia begins to influence substance use during emerging adulthood, suggesting that studying this developmental period may be critical to disentangling the complex relationship between these two phenotypes.

Interestingly, polygenic scores for schizophrenia did not predict phenotypes that only involved alcohol consumption: alcohol use and binge drinking. In contrast, a small genetic correlation between schizophrenia and alcohol consumption was recently reported in a sample of older adults (Clarke *et al*., 2017). Given the relatively weak genetic correlation between schizophrenia and alcohol consumption (*r_g_* = .13), it is possible that we were not powered to detect cross-trait effects. Alternatively, different genetic factors may influence alcohol consumption at different stages of development (Edwards and Kendler, 2013). In the present sample of emerging adults, alcohol use and binge drinking are relatively normative behaviors and, as such, they may be less influenced by genetic factors during this developmental period. Indeed, research has demonstrated that genetic influences on alcohol consumption typically increase across the lifespan (Kendler *et al*., 2008; van Beek *et al*., 2012).

Our findings should be interpreted in light of several limitations. First, our measure of illicit drug use did not identify the specific substance that was consumed, so we have limited insight into substance-specific patterns of drug use. However, the monthly rates of alcohol and drug use observed in this study are quite similar to those reported by college students in the *Monitoring the Future* study (Johnston *et al*., 2011), so we are still able to generate insight into general patterns of substance use. Second, our analyses were restricted to non-Hispanic European participants to reduce the risk of spurious findings caused by population stratification. Consequently, the findings of our study may not generalize to other ancestral populations. A third potential limitation is our relatively moderate sample size. However, concerns about statistical power in the present study are partially attenuated by the fact that (i) we were well-powered for our within-person approach, (ii) we leveraged *a priori* effect size estimates from a well-powered GWAS of schizophrenia (Schizophrenia Working Group of the Psychiatric Genomics Consortium, 2014), and (iii) we examined aggregate genomic variation rather than individual SNPs of small effect.

Despite these limitations, this study exhibited notable strengths in its novel approach. As the first longitudinal, event-level investigation of genetic risk for schizophrenia and its effects on substance use in daily life, this work provides ecologically valid evidence that these two psychiatric conditions are influenced, in part, by shared genetic factors. Notably, this study demonstrated that genetic risk for schizophrenia can predict important behavioral phenotypes in a sample of healthy, university students, where schizophrenia prevalence is expected to be minimal. In doing so, we present a critical extension of previous work, which has primarily examined the genetic underpinnings of substance use in clinical samples. Moreover, the present study contributes to the broader literature by illustrating how a novel combination of polygenic prediction and intensive longitudinal methods can be used to characterize broad and developmentally-specific effects of genetic variation.

As repeated event-level measurement can be used to examine concurrent dynamic processes, person-centered approaches may facilitate greater insight into the specific temporal dynamics or causal relationships of co-occurring phenomena. For instance, future studies that employ similar methods may be uniquely poised to elucidate *how* genetic risk for schizophrenia influences cannabis use during emerging adulthood by simultaneously assessing daily experiences with prodromal symptoms and cannabis use. As interest in the study of genetic contributions to psychiatric outcomes continues to rise, it will be important to use person-centered approaches that characterize the relationship between related phenomena as it unfolds in the life of an individual person. In doing so, future studies using polygenic prediction methods will be better suited to investigate *when* and *how* those predispositions contribute to complex human behavior.

